# Physiological response of *Microcystis aeruginosa* exposed to aqueous extracts of *Pistia stratiotes* and *Pontederia crassipes*

**DOI:** 10.1101/2024.09.30.615933

**Authors:** Luan de Oliveira Silva, Allan Amorim Santos, Evelyn Maribel Condori Peñaloza, Ana Beatriz Furlanetto Pacheco, Sandra Maria Feliciano de Oliveira e Azevedo

## Abstract

Macrophyte extracts inhibit cyanobacteria growth, offering a sustainable solution for bloom control. The present study aimed to evaluate *Microcystis* aeruginosa’s response to aqueous extracts obtained from the dried biomass of *Pistia stratiotes* L. and *Pontederia crassipes* Mart. Solms. Physiological parameters analyzed included growth, photosynthesis, and antioxidative response. Reactive oxygen species generation and the chemical profile of the aqueous extracts were analyzed. At 4.0 g L^-1^, both extracts inhibited cyanobacterial growth in 6 days. *P. stratiotes* extract achieved 100% inhibition, while *P. crassipes* extract reached 60%. *P. stratiotes* extract showed higher photosynthetic inhibition (99% vs. 12% for *P. crassipes*). This was related to the downregulation of the *psbA* gene (coding for the photosystem II protein D1). Exposure to both extracts increased intracellular reactive oxygen species content in cyanobacterial cells and increased peroxiredoxin gene (*prxA)* transcripts. This response was not noted for *sod* (superoxide dismutase) transcripts, although SOD enzymatic activity increased in cultures exposed to *P. stratiotes* extracts. Upon incubation of the macrophyte extracts with *M. aeruginosa* cultures, phenol concentrations decreased, and their general metabolic profile changed. Thus, *P. stratiotes* extract outperformed *P. crassipes*, inhibiting *M. aeruginosa* growth, affecting photosystem II, and inducing oxidative stress.

**Highlights:**

- Aqueous extracts from dried biomass of *P. stratiotes* and *P. crassipes* inhibited *M. aeruginosa* growth.
- *P. stratiotes* extract suppressed photosystem II activity, while *P. crassipes* did not
- Both extracts elicited ROS production in cells.
- Peroxiredoxin gene expression upregulated by extract exposure.
- *P. stratiotes* extract increased SOD activity in cells.

## 1 INTRODUCTION

Water quality is deteriorating worldwide due to human impacts, such as urban and industrial effluents being discharged into aquatic ecosystems, which accelerates eutrophication (Le Moal et al., 2019). One consequence of eutrophication is the development of harmful algae blooms (HABs), especially those formed by cyanobacteria (Paerl and Barnard 2020). It is known that several cyanobacteria species can produce toxic metabolites (cyanotoxins), which affect aquatic biota and human health (Buratti et al., 2017). The most common bloom-forming cyanobacteria in freshwater is *Microcystis aeruginosa*. In temperate climates, *Microcystis* cells grow in the water column during spring, and blooms develop in summer, whereas in the colder seasons, colonies move towards the sediment/bottom. High temperatures and high sunlight irradiance in tropical regions contribute to perennial year-round blooms (Vanderley et al., 2021). The concern about these blooms’ occurrence is mainly related to the capability of some strains to produce microcystin, a potent cyanotoxin involved in cases of animal and human poisoning (Azevedo et al., 2002; Harke et al., 2016; Polyak and Polyak 2022).

Considering the negative effects of HABs, different methods have been proposed to mitigate these events, including chemical, physical, or biological interventions, with limited success in field application. A recent meta-analysis concluded that only some chemical methods (e.g. the addition of copper sulfate or hydrogen peroxide to the water) improved water quality in field conditions (Anantapantula and Wilson 2023). However, these substances may negatively affect non-target organisms, unbalancing the aquatic food web (Silva et al., 2018; Santos et al., 2021).

Biological strategies for bloom control include the biomanipulation of biotic interactions involving fish, zooplankton, and macrophytes (Triest et al., 2016; Mohamed 2017; Pal et al., 2020; Ly et al., 2024). Most studies using macrophytes explored submerged species that promote a clear-water state mediated by competition with phytoplankton for nutrients, long zooplankton permanence, and allelopathic inhibition of phytoplankton growth (Triest et al., 2016). Floating macrophytes such as *Pistia stratiotes, Salvinia* sp., and *Pontederia crassipes*, also present these advantages and, additionally, their growth limit light incidence in the water column (Srivastava et al., 2017; Jinqing Wang et al., 2018).

*P. stratiotes* and *P. crassipes,* popularly known as water lettuce and water hyacinth, respectively, are cosmopolitan species widely spread in eutrophic environments (Galal et al., 2019; Datta et al., 2021). In these conditions, these macrophytes can grow to form dense mats, causing economic and ecological problems (Datta et al., 2021). This situation requires the continuous removal of the plants, forming large amounts of biomass. Although this residual biomass may be useful as fertilizer or as a substrate for constructed floating wetlands (small artificial platforms that allow aquatic plants to grow and remove nutrients and other molecules from the water column (Kochi et al., 2020; Poveda 2022), the amount removed from the water usually exceeds its use and generates an environmental hazard.

Besides, it is already known that many species of macrophytes release allelochemicals that suppress cyanobacterial growth (Mohamed 2017; Amorim et al., 2019; De Lima et al., 2023). Diverse chemicals mediate allelopathy; for example, polyphenols, alkaloids, and nonanoic and succinic acids are known to inhibit the growth of cyanobacteria (Chen et al., 2021; Li et al., 2016; Yotsova et al., 2017; Zhang et al., 2010; Zhao et al., 2020). The underlying mechanisms of action are related to the decrease of chlorophyll-*a* (Chl-*a*) concentration, inhibition of photosynthesis, and promotion of oxidative stress (Liu et al., 2015; Qian et al., 2010; X. X. Wu et al., 2019). However, most studies that characterized inhibition routes evaluated the effects of isolated allelochemicals and did not consider the complex matrices of chemical compounds released by macrophytes into aquatic environments.

The allelochemical potential of aquatic plants can be managed as a nature-based solution to control harmful cyanobacteria since it is an effective, economical, and ecologically safe alternative due to its high inhibitory rate, high availability, and low cost (Mohamed 2017). The allelopathic effect of macrophytes *Pistia stratiotes* and *Pontederia crassipes* on *M. aeruginosa* has been explored using plant tissues, exudates, or purified compounds (X. Wu et al., 2013, 2015; X. X. Wu et al., 2019; Lourenção et al., 2021). Interestingly, the potential use of aqueous extract from macrophytes, which is much closer to the environmental scenario than solvent extracts or purified products, has been poorly investigated. It is a basic step to explore alternative uses for the residual biomasses, which includes a nature-based solution for mitigating cyanobacteria and improving the water quality. Therefore, the main goal of this study was to test the effect of aqueous extracts obtained from the dry biomass of *P. stratiotes* and *P. crassipes* on the physiology of *M. aeruginosa*, exploring the growth, photosynthetic activity, and oxidative response of cyanobacterial cells, contributing to understanding the underlying mechanisms of the allelopathic effects of these macrophytes on cyanobacteria.

## 2 MATERIALS AND METHODS

### 2.1 Cyanobacterial and macrophyte culture conditions

A *Microcystis aeruginosa* strain (LETC-MC-32) isolated from a coastal lagoon (Jacarepaguá lagoon, 22°59’00.4” S 43°24’36.2” W) in Rio de Janeiro, RJ (Brazil) was used. The cells were cultivated in ASM-1 (Gorham et al., 1964) with a light intensity of 30 µmol photons m^-2^ s^-1^, 12h light: dark cycle and 25 ± 1 °C temperature.

The macrophytes used were *P. stratiotes* L. and *P. crassipes* Solms. Specimens were collected in September 2018 in a hydroelectric power plant reservoir (Barra do Braúna, 42°24′26, 4” W 21°26’51, 5” S) located in the Pomba River, Minas Gerais State, Brazil. The macrophytes were taken to the laboratory in plastic bags containing reservoir water. Then, they were maintained in plastic boxes filled with 40 L of distilled water supplemented with nutrients (NPK 10:10:10). The macrophytes were maintained at 24 ± 2 °C, 1000 μmol photons m^−2^ s^−1^, and a 12h light: dark cycle.

### 2.2 Preparation of *P. stratiotes* and *P. crassipes* aqueous extracts

Adult plants of similar weight were washed with potable water for each macrophyte species and dried in a forced-air oven at 60 °C until they reached a constant weight. The resulting material was grounded in a mill to obtain a powder. Approximately 96 g of powder was homogenized in deionized water (6 L) over three h at 30 °C. The resulting aqueous extracts (16.0 g L^-1^) were filtered through 0.45 µm pore membranes, and the resulting filtrates were stored at -20 °C in 200 mL aliquots. Before the experiments, extract aliquots were thawed and added with nutrients to reproduce the composition of the ASM-1 medium. The resulting media were filtered through a 0.22 µm pore membrane to remove suspended particles and to avoid microbial contamination.

### 2.3 Experimental setup

Initially, an experiment was carried out to evaluate which concentration of the macrophyte aqueous extracts would inhibit *M. aeruginosa* growth. From the original aqueous extract (16.0 g L^-1^), dilutions were made in ASM-1 to reach the concentrations 0.5, 1.0, 2.0 and 4.0 g L^-1^. A culture of *M. aeruginosa* in the exponential phase was used as an inoculum to initiate cultures with 1.0 x 10^5^ cells mL^-1^. The treatments consisted of cultivating *M. aeruginosa* in ASM-1 medium prepared with aqueous extracts of *P. stratiotes* or *P. crassipes*. The control consisted of cultivating *M. aeruginosa* in the original ASM-1 medium. Cultures (120 mL) were maintained for ten days at 30 µmol photons m^-2^ s^-1^, 12h light: dark cycle, temperature of 25 ± 1 °C. The experiments were conducted in Erlenmeyer flasks with four replicates for each condition. Sampling was taken every three days to measure of Chl-*a* concentration and estimate cyanobacterial growth (Table 1).

**Table 1.**
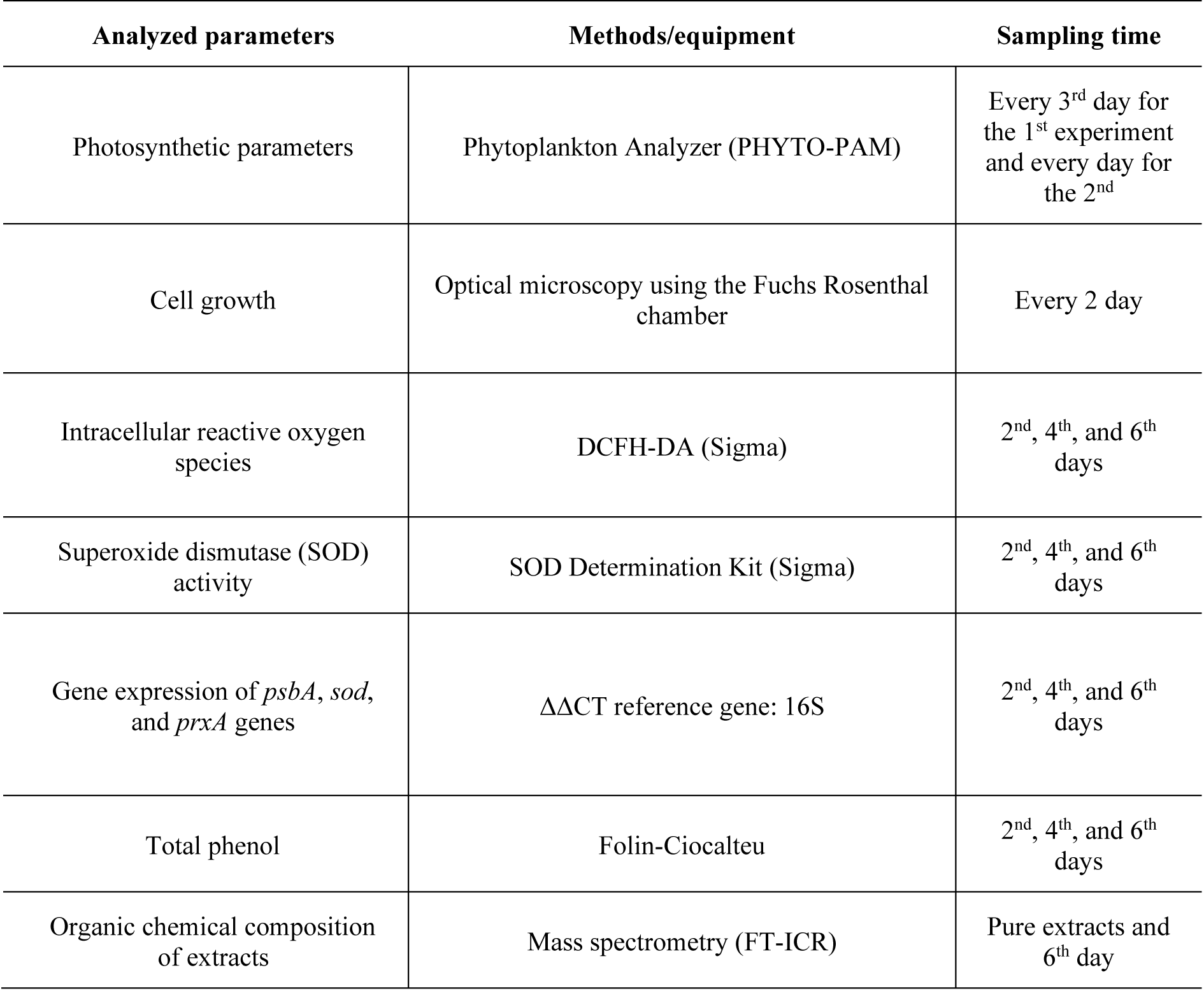
Description of the analytical methods and their respective sampling time for analysis.

In a subsequent experiment, a culture of *M. aeruginosa* in the exponential phase was used as an inoculum to initiate cultures with 5.0 x 10^5^ cells mL^-1^. The treatments consisted of cultivating *M. aeruginosa* in an ASM-1 medium prepared with aqueous extracts of 4.0 g L^-1^ of *P. stratiotes* or *P. crassipes*. The control consisted of the cultivation of *M. aeruginosa* in ASM-1 medium. Cultures (1 L) were maintained in Erlenmeyer flasks (n=4) for eight days at 30 µmol photons m^-2^ s^-1^, 12h light: dark cycle, temperature of 25 ± 1 °C. Sampling was done every day for the determination of Chl-*a* concentration and photosynthetic activity, every two days for cell counting and total phenol quantification, on days 2, 4, and 6 for RNA extraction and reactive oxygen species (ROS) estimation, and days 0 and 6 for the chemical characterization of the extracts (Table 1).

### 2.4 Cyanobacterial growth and photosynthetic activity

The cell density of *M. aeruginosa* cultures was monitored by optical microscopy using a Fuchs-Rosenthal chamber. Values were expressed as cell mL^-1^, and the inhibition rate was calculated as follows:

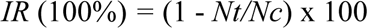

Where *IR* means inhibition rate; *Nt* and *Nc* indicate the number of cells in treatment (cultivation with macrophyte extracts) and control (ASM-1), respectively.

Photosynthetic activity was measured using a PHYTO-PAM fluorometer (Heinz Walz GmbH, Germany) equipped with a PHYTO-EDF detection unit for measuring cyanobacteria fluorescence. Saturation pulses (36 µmol photons m^-2^/s^-1^) were applied, and fluorescence data were converted to Chl-*a* concentration (μg L^-1^) and photosynthetic yield (relative Fv/Fm) related to the photosystem II.

### 2.5 Determination of reactive oxygen species (ROS) production

A volume of 40 mL of cyanobacteria culture was centrifugated (12000 x g for 10 minutes at 4 °C), and the cells were resuspended in appropriate ASM-1 volumes to obtain the same cell density (1.0 x 10^7^ cells mL^−1^).

Intracellular ROS production by *M. aeruginosa* was measured by adding 10 µM (final concentration) of DCFH-DA in DMSO to 500 µL of cell culture and incubating on a shaker at room temperature in the dark. After 30 min, samples were subjected to fluorescence spectrophotometric analysis. DCFH-DA is a nonpolar dye that is converted into the polar derivative DCFH by cellular esterases. DCFH is nonfluorescent but switches to highly fluorescent DCF when oxidized by intracellular ROS or other peroxides, with an excitation wavelength of 485 nm and an emission band between 500 and 600 nm.

### 2.6 Superoxide dismutase activity

A 120 mL volume of cyanobacteria culture was centrifugated (12000 x g for 10 minutes at 4 °C), and the cell pellet was resuspended in ASM-1 (1 mL). The cell suspensions were transferred to a 2 mL Lysis Matrix Tube with glass beads (150-212 µm, glass beads acid washed, Sigma) and submitted to lysis in FastPrep™ equipment (6 cycles of 20 sec, 4.5 m s^-1^). After bead decantation, the homogenate was transferred to a new tube and centrifuged (15000 g for 1 min at 4 °C). The resulting supernatant (cell lysate) was used in the assay. The protein concentrations were determined in the resulting lysates using the Qubit™ Protein and Protein Broad Range (BR) Assay Kit and a Qubit (Thermo Fisher Scientific) fluorimeter.

The activity of the superoxide dismutase (SOD) enzyme was measured in cell lysates using the SOD Determination Kit (Sigma Merck®), and calculations were carried out following the manufacturer’s instructions. The values obtained for enzymatic activity were normalized using protein concentrations. Results were represented as a ratio between SOD activity obtained for treatment (cultures exposed to aqueous extracts) samples and SOD activity obtained for the control (cultures in ASM-1).

### 2.7 Analysis of the Relative Transcript Abundance

Transcript levels of the *psbA*, *sod*, and *prxA* genes were measured. Cyanobacterial cells (80 mL of culture) were concentrated by centrifugation (15000 x g for 15 minutes at 4 °C), and pellets were resuspended in 300 μL of ASM-1 and immediately frozen in liquid nitrogen.

RNA extraction was performed using the NucleoSpin extraction kit (MACHEREY-NAGEL) following the manufacturer’s instructions. RNA was quantified in Nanodrop (Thermo Fisher Scientific) and immediately converted into cDNA. According to the manufacturer’s instructions, the cDNA synthesis was performed with the enzyme GoScript reverse transcriptase (Promega) and random primers. cDNA was diluted (1:10 and 1:100) and stored at -80 °C.

The cDNAs obtained were used in qPCR. In each reaction, 3 µL of cDNA (1:100 dilution for the *psbA* gene; 1:10 dilution for the other genes) were added to 7.5 µL of Sybr Green PCR MasterMix (Thermo Fisher Scientific), 0.6 µL of each primer (20 µM) and nuclease-free water to a total volume of 15 µL. Reactions were performed in triplicate in a QuantStudio 3 Real-Time PCR (Thermo Fisher Scientific) equipment with incubation at 95 °C for 10 min followed by 40 cycles of 95 °C for 15 s + 60 °C for 1 min (Jia Wang et al., 2018). The 16SrRNA was amplified in parallel and used for normalization of target transcript abundances (Jia Wang et al., 2018). The relative abundance of transcripts was estimated according to ΔΔCT (Livak and Schmittgen 2001), considering the control as the normalizing condition for each sampling time. The primers used are listed in Supplementary Table 1.

### 2.8 Quantification of phenolic compounds

Aliquots of 1 mL of each macrophyte aqueous extract were filtered in 0.22 µm pore membranes, and the filtered volume was used to determine the concentration of total phenols by the Folin-Ciocalteu method (Box 1983). This method was adapted to microplates with a final reaction volume of 300 µL, consisting of 240 µL filtered sample, 40 µL of Na_2_CO_3_ (200 g L^-1^) in 0.1 N NaOH, and 20 µL Folin-Ciocalteu reagent. Samples were kept in the dark for 20 min, and the absorbance was measured in a spectrophotometer at a wavelength of 630 nm. The concentration of phenols in the samples was calculated using a standard calibration curve prepared with gallic acid.

### 2.9 Mass spectrometry (FT-ICR-MS)

The analysis was carried out to compare the chemical profile of (i) the original aqueous extracts of *P. stratiotes* and *P. crassipes* diluted at 4 g L^-1^ (in the base of the results of the experiment 2.3) codified as T0PC and T0PS, (ii) extracts obtained after 6 days of incubation in cultures of *M. aeruginosa* codified as T6PC and T6PS, and (iii) extracts obtained after 6 days of incubation, without the presence of cyanobacteria codified as POSPC and POSPS. Extracts were maintained under the same conditions described in 2.3. Aliquots of 40 mL were obtained, lyophilized (4 replicates for each condition), and the material was resuspended in water/acetonitrile 1:1 at a final concentration of 1 mg mL^-1^ and filtered through 0.22 μm pore membranes.

Analysis was performed on a Bruker SolariX XR, 7 Tesla Fourier Transform Ion Cyclotron (FT-ICR-MS) instrument (Brüker Daltonics Inc., Billerica, MA) equipped with an electrospray ionization (ESI) source. Samples were infused directly at a flow rate of 3.0 µL min^-1^. The ESI source parameters in positive and negative ionization modes were: nebulizer gas pressure of 1.0 bar, temperature of 200 °C, dry gas of 4 L min^-1^ and capillary potential of 5.0 kV. Spectra were acquired with a mass range of m/z 75.26 to 1200 (with a resolution of 200 000 at m/z 400), 10 accumulated scans for each sample, and mass accuracy of < 2 ppm, provided for molecular formula assignments unambiguous for a single charge of molecular ions.

The data processing was carried out using the mMass software (Open Source Mass Spectrometry Tool, version 5.5.0). Data analysis was normalized using the MetaboAnalyst platform 4.0 (CHONG et al.,, 2019). Principal Component Analysis (PCA) was performed using the MetaboAnalyst 4.0 platform to distinguish the effect of time and experimental conditions. For this purpose, a data matrix for ESI (-) and ESI (+) ionization modes was generated (m/z variables detected from 75.26 to 1200) for the macrophyte in the following conditions: i) extracts of *P. stratiotes* and *P. crassipes* incubated separately for 6 days or ii) maintained in *M. aeruginosa* cultures for 6 days (filtered with a 0.22 μm pore) and iii) the aqueous extracts on time 0.

### 2.10 Statistical analyses

Cell density, Chl-*a*, and photosynthetic efficiency (yield) were evaluated using Two-way analysis of variance (ANOVA) following the required assumptions. For the other tests, a one-way analysis of variance (ANOVA) was performed. When significant differences were detected in ANOVA tests, Tukey’s HSD post hoc test was used to separate the means. Comparisons were performed within each sample time, and the differences were carried out at a 5% significance level using GraphPad Prism 8.0 software.

## 3 RESULTS

### 3.1 Effects of macrophyte aqueous extracts on cyanobacterial growth

The aqueous extract of *P. stratiotes* at concentrations 1.0, 2.0, and 4.0 g L^-1^ significantly reduced *M. aeruginosa* growth (estimated by Chl-*a* concentration) compared to the control condition over time (days 3 to 10) (Figure 1A). The maximum inhibition with 4.0 g L^-1^ on days 6 and 10 when the Chl-*a* concentration reduced by 95% compared to the control. The cultivation of *M. aeruginosa* in the presence of the aqueous extract of *P. crassipes* resulted in a significant growth reduction at the highest concentration of dry biomass tested (Figure 1B) (p < 0.05). On day 10, Chl-*a* concentrations in cultures with *P. crassipes* extracts were approximately 30% of those maintained in control conditions. Based on these results, for both *P. stratiotes* and *P. crassipes*, aqueous extracts with a concentration of 4.0 g L^-1^ of macrophyte dry biomass were used in subsequent experiments.

**Fig. 1.**
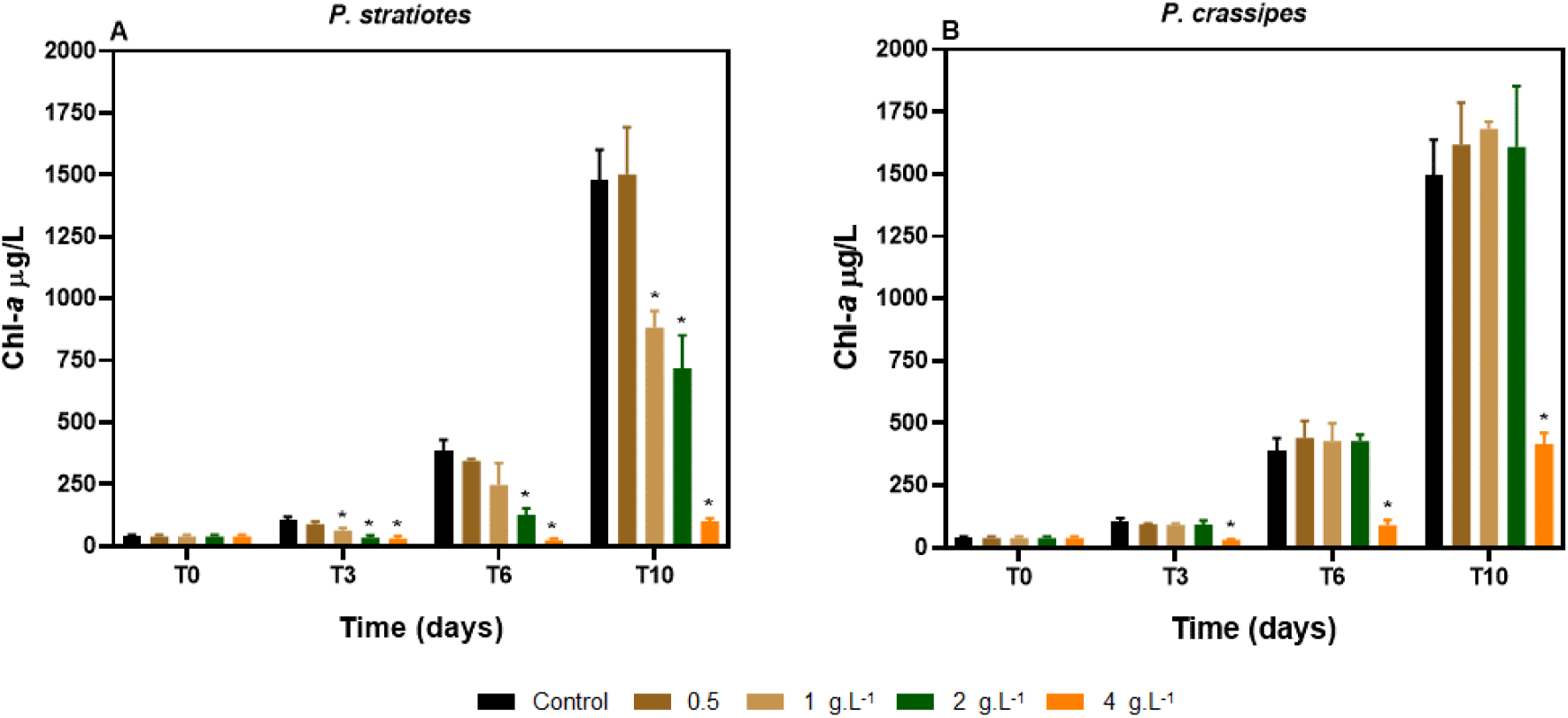
Effects of different concentrations of macrophyte aqueous extracts on *M. aeruginosa* growth. Cyanobacterial growth was followed by Chl-*a* concentration for 10 days. (**A**) extract from *P. stratiotes* and (**B**) extract from *P. crassipes*. Significant differences between control and treatment within each sampling time are indicated (*) (p<0.05).

For the next experiments, both macrophyte extracts were used at a concentration of 4.0 g L^-1^, as it caused maximum inhibition of cyanobacterial growth for 10 days. Growth of *M. aeruginosa* cultures added with *P. stratiotes* or *P. crassipes* aqueous extracts was monitored by cell density for 8 days and compared to control conditions (ASM-1). For both aqueous extracts, there was a significant reduction in cyanobacterial cell density (p < 0.05) from day 2^nd^ to day 8^th^ (Figure 2A). The aqueous extract of *P. stratiotes* reduced the growth of *M. aeruginosa* to a greater extent than that of *P. crassipes*. This was represented as a growth inhibition ratio, showing that although both extracts inhibited 50% of cyanobacteria growth on the 2^nd^ day, the *P. stratiotes* extract caused an inhibition of 95% on the 4^th^ day and 100% afterward (Figure 2B). The presence of the *P. crassipes* extract caused a smaller inhibitory effect, a maximum of 60-70%, which still allowed cyanobacterial growth.

**Fig. 2.**
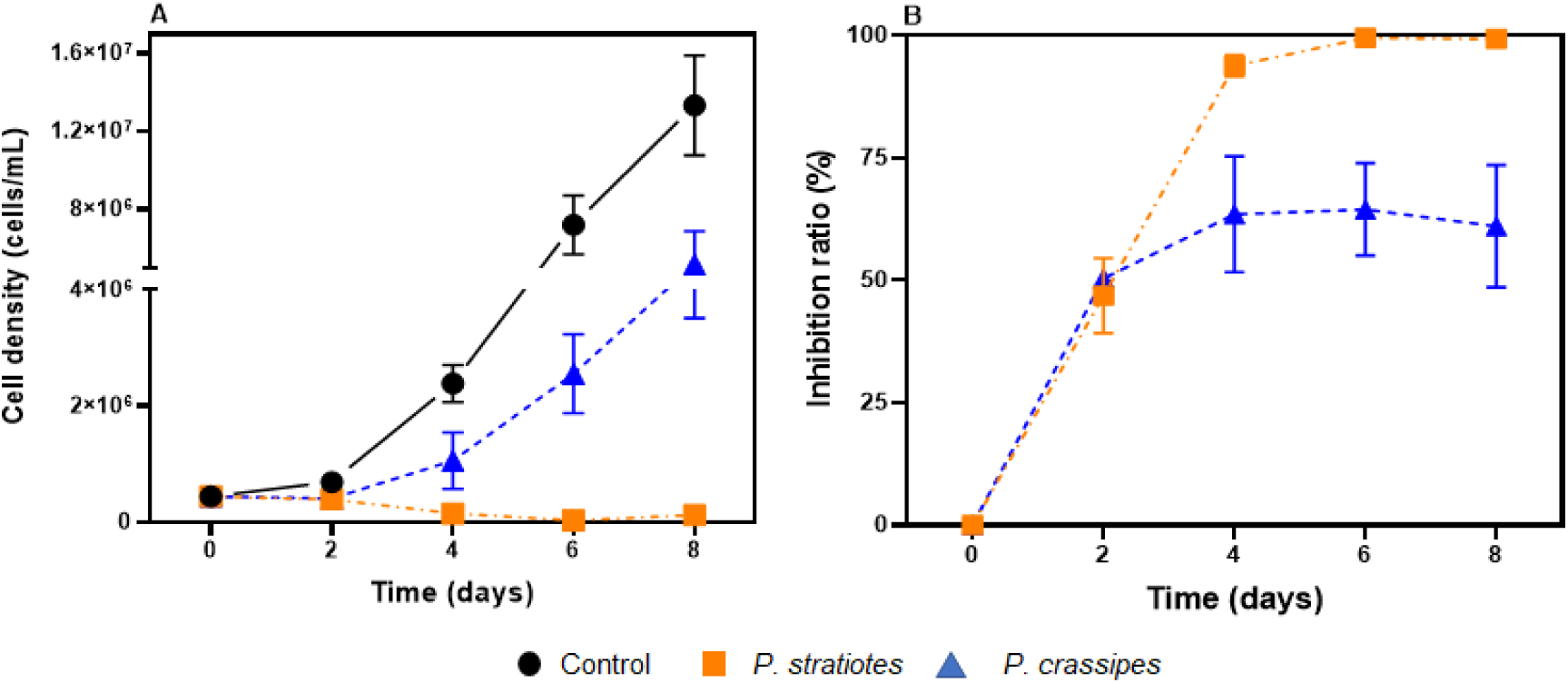
Effects of *P. stratiotes* and *P. crassipes* aqueous extracts (4 g L^-1^) on *M. aeruginosa* growth. (A) cell density and (B) inhibition ratio, representing growth inhibition compared to control.

### 3.2 Effects of macrophyte extracts on cyanobacterial photosynthesis

Cultures of *M. aeruginosa* maintained with *P. stratiotes* or *P. crassipes* aqueous extracts (4.0 g L^-1^) showed lower Chl-*a* concentrations than the control from day 2 (p < 0.05) (Figure 3A). From day 3^rd^, a stronger effect was noted for cyanobacteria cultures incubated with the *P. stratiotes* extract than for those incubated with *P. crassipes* extract (p < 0.05). On the 8^th^ day, Chl-*a* concentrations represented 10% of the control for cultures with *P. stratiotes* extracts, and 50% of the control for those with *P. crassipes* extracts.

**Fig. 3.**
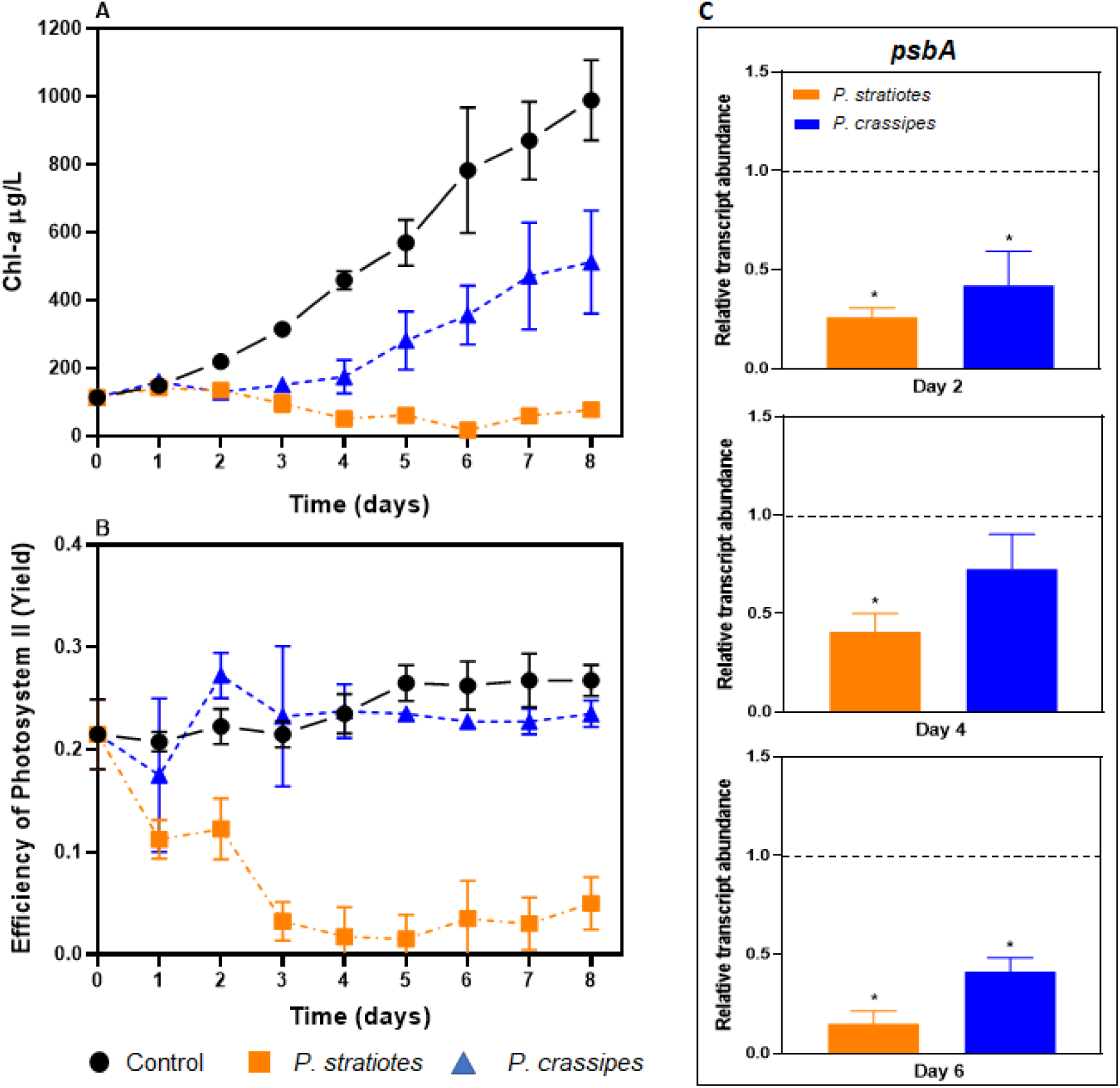
Effects of *P. stratiotes* and *P. crassipes* aqueous extracts (4 g L^-1^) on *M. aeruginosa* photosynthetic parameters. (A) Chl-*a* concentration; (B) efficiency of photosystem II; (C) relative abundance of *psbA* gene transcripts. The dashed line represents control. Significant differences between control and treatment within each sampling time are indicated (*) (p<0.05).

The decrease in Chl-*a* concentration was reflected in the photosynthetic efficiency of photosystem II (PSII) in *M. aeruginosa* cultures maintained in *P. stratiotes* extracts (Figure 3B). The extract had a significant inhibitory effect (p < 0.05) compared both to the control and the treatment with the *P. crassipes* from day 1^st^ on. There were no significant differences between the treatment with *P. crassipes* extract and the control, indicating that the inhibitory effect of this extract on cyanobacteria growth may not be related to PSII damage.

We also investigated the potential impact of the exposure to macrophyte extracts on the synthesis of the D1 protein in *M. aeruginosa* cells, estimating the relative abundance of the corresponding gene (*psbA*) transcripts. Compared to the control, incubation with *P. stratiotes* extracts resulted in a decrease in *M. aeruginosa psbA* transcripts at all tested times, days 2, 4, and 6 (p < 0.05) (Figure 3C), while for cultures maintained with *P. crassipes* extracts decreases were observed on day 2 and 6.

### 3.3 Effects of macrophyte extracts on the cyanobacterial anti-oxidant response

When *M. aeruginosa* cells were exposed to macrophyte aqueous extracts, intracellular ROS contents increased significantly over time compared to control (p < 0.05), with similar rises for both macrophyte extracts (Table 2). On day 2, ROS concentrations in *M. aeruginosa* cells cultivated with the extracts were approximately 10 times higher than the control and increased to values 40 times higher after 6 days.

**Table 2.** Effects of *P. stratiotes* and *P. crassipes* aqueous extracts (4.0 g L^-1^) on ROS production in *M. aeruginosa* cultures. Values represent the average (std ± dev) of fluorescence intensities and fold change represents how much ROS increased compared to the control condition.

| Samples | Ratio ± standard<br>deviation | Ratio ± standard<br>deviation | Ratio ± standard<br>deviation |
| --- | --- | --- | --- |
|  | T=2 | T=4 | T=6 |
| Control | 7.90 ± 0.05 | 6.37 ± 1.20 | 10.73 ± 1.15 |
| *P. stratiotes* | 96.91 ± 4.27 | 264.05 ± 56.77 | 365.32 ± 126.33 |
| Fold change | 12.09 ± 0.54 | 31.57 ± 7.18 | 46.80 ± 16.10 |
| *P. crassipes* | 58.56 ± 13.02 | 394.13 ± 121.91 | 428.47 ± 235.20 |
| Fold change | 8.06 ± 1.64 | 47.57 ± 15.43 | 44.23 ± 29.77 |

The relative abundance of the *prxA* and *sod* gene transcripts and the SOD activity was estimated as an approximation of the anti-oxidative response of *M. aeruginosa* cells (Figure 4). Compared to the control condition, there was a significant upregulation of the *prxA* gene (p < 0.05) on the 2^nd^ day of exposure to the macrophyte extracts, with a more prominent effect for the *P. crassipes* extract treatment, but only for the *P. stratiotes* extract this increase remained on the 4^th^ day (Figure 4A). On the 6^th^ day, *prxA* transcript abundance decreased in cells treated with the *P. crassipes* extract compared to the control.

**Fig. 4.**
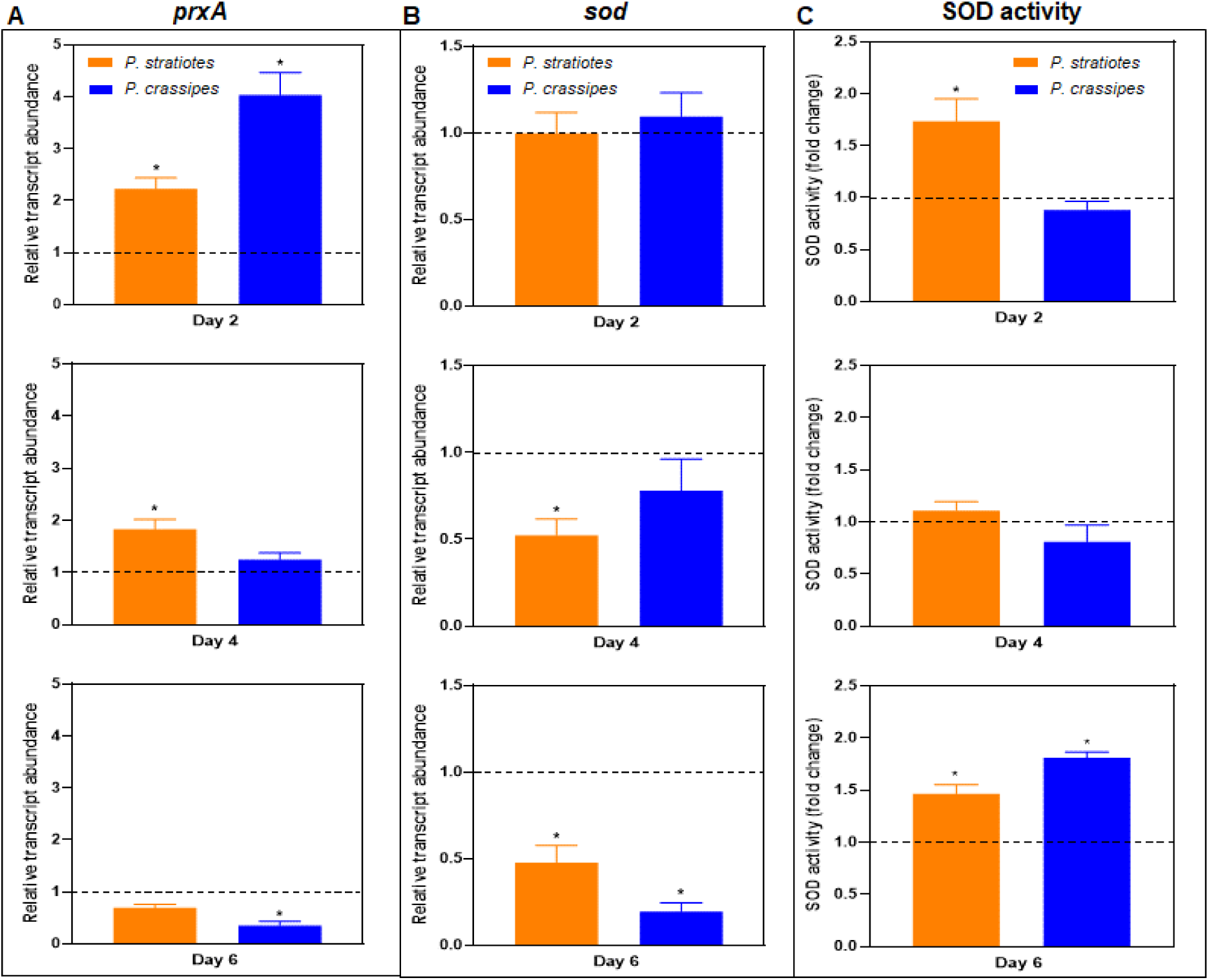
Effects of *P. stratiotes* and *P. crassipes* aqueous extracts (4 g L^-1^) on *M. aeruginosa* anti-oxidant response. (A) relative abundance of *prxA* transcripts; (B) relative abundance of *sod* transcripts; (C) superoxide dismutase activity. All values are represented as fold change in comparison to the control condition. The dashed line represents control. Significant differences between control and treatment within each sampling time are indicated (*) (p<0.05).

There was no significant difference in *sod* transcript abundances on the 2^nd^ day of exposure to macrophyte extracts (Figure 4B). In subsequent samples, *sod* transcripts were less abundant than in the control (p < 0.05). SOD enzymatic activity was higher on day 2 in cells treated with the *P. stratiotes* extract (p < 0.05) (Figure 4C). After 4 days, there was no significant difference comparing each treatment to the control, but on the 6^th^ day, SOD activity was higher in the treatments than in the control.

### 3.4 Chemical composition of macrophyte aqueous extracts

The original concentrations of total phenols in the macrophyte aqueous extracts were 17 mg L^-1^ and 12 mg L^-1^ for *P. stratiotes* and *P. crassipes*, respectively (Figure 5). Incubation of *M. aeruginosa* cells with the aqueous extracts led to a decrease in these values (p < 0.05), particularly for *P. crassipes* extracts. In contrast, when the aqueous extracts were incubated for 6 days without *M. aeruginosa* cultures, the phenol concentrations remained constant.

**Fig. 5.**
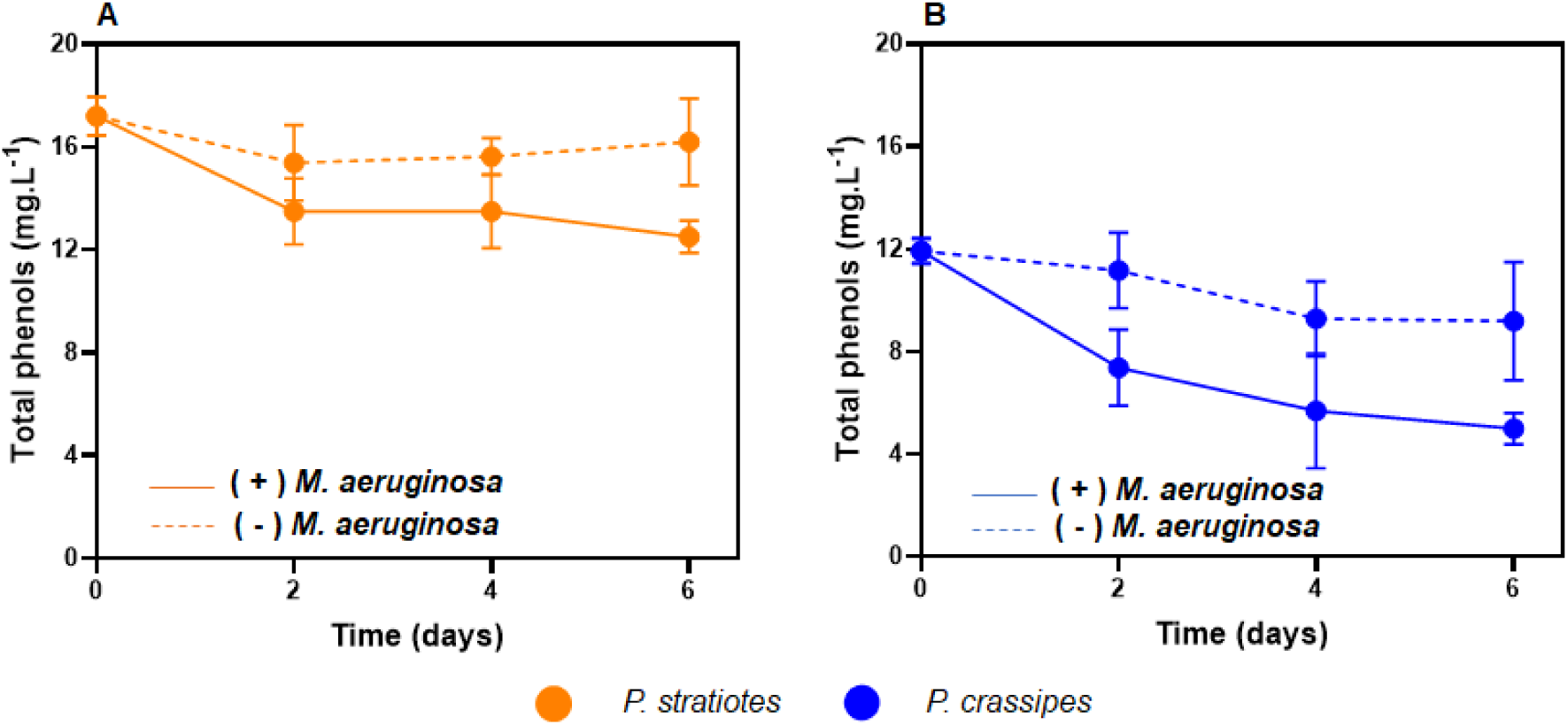
Total phenol concentrations in macrophyte aqueous extracts. (A) *P. stratiotes* aqueous extracts incubated with (+) or without (-) *M. aeruginosa*; (B) *P. crassipes* aqueous extracts incubated with or without *M. aeruginosa.* A one-way ANOVA test was performed, and differences over time were analyzed.

A partial chemical characterization of aqueous extract from *P. crassipes* and *P. stratiotes* was performed by analyzing FT-ICR-MS data. For each macrophyte, the original extract (T0) was compared with extracts maintained for 6 days with the presence of *M. aeruginosa* (T6PS and T6PC) and with extracts maintained for 6 days without the presence of *M. aeruginosa* (POSPC and POSPS). PCA was performed to explore and interpret those FT-ICR-MS data wherein the scores plot of PC1 and PC2 allowed the discrimination of the three conditions analyzed for each macrophyte (Figure 6A and D). The loadings plot with the highest values showed that the peaks at *m/z* 317.11557 from T6PC samples (Figures 6B and C) and the peak *m/z* 814.58053 from T6PS samples (Figures 6E and F) contribute significantly to the differentiation of the samples. However, these peaks were not prominent in respective macrophyte extracts from T0 or those incubated without *M. aeruginosa* (POSPC and POSPS). Thus, these molecules could be related to the metabolic activity of *M. aeruginosa* cultures.

**Fig. 6.**
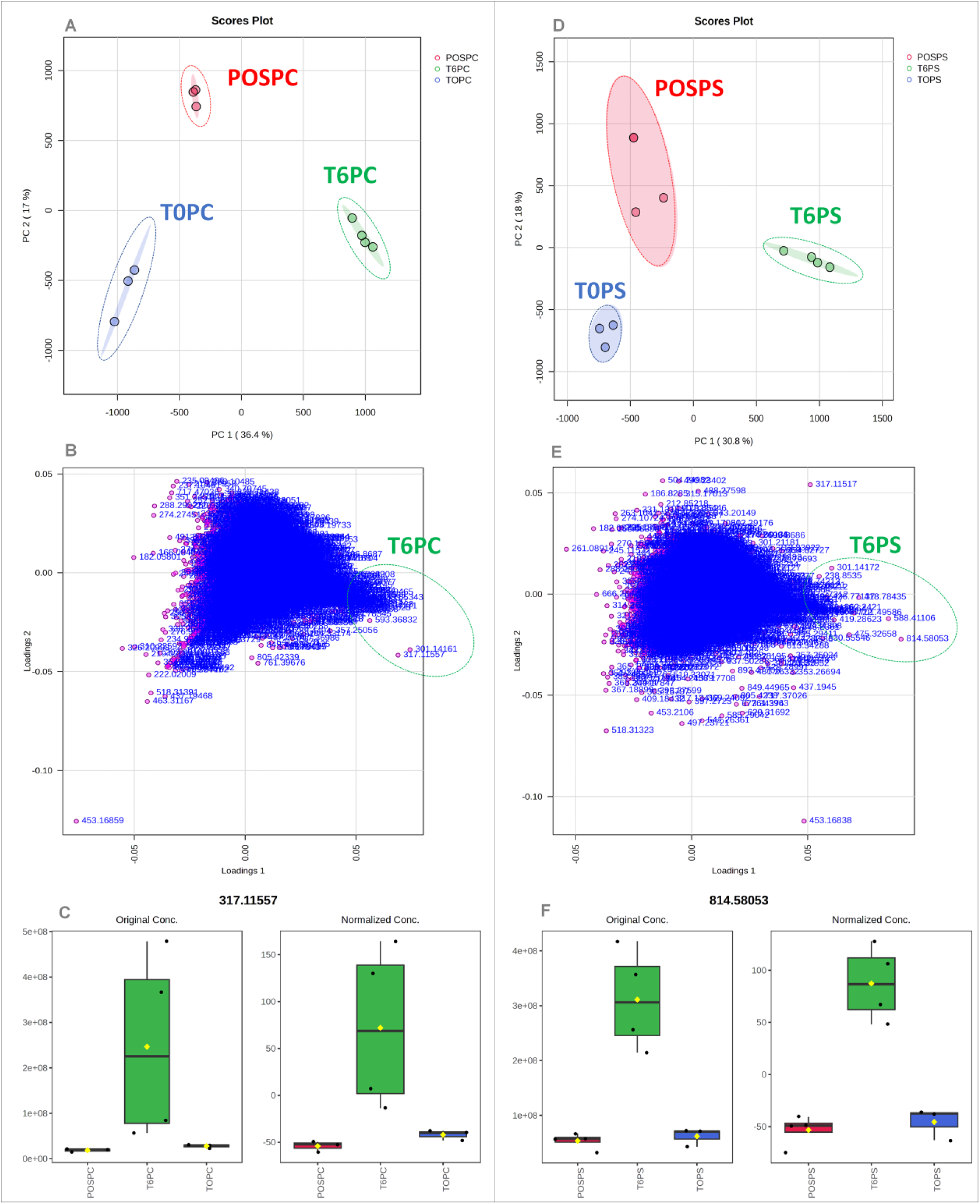
Principal component analysis (PCA) is based on the metabolic composition of aqueous extracts of *P. stratiotes* and *P. crassipes* using mass spectrometry in positive mode. T0PS and T0PC mean original extracts of *P. crassipes* and *P. stratiotes*, respectively, time 0; T6PS and T6PCmean extracts after 6 days of incubation with the presence of *M. aeruginosa,* and POSPC and POSPS mean extracts after 6 days of incubation without the presence of *M. aeruginosa*. (A and B) score and loading of *P. crassipes* extracts; (C) *m/z* 317.11557 feature with highest loading representing the group T6PC. (D and E) score and loading of *P. stratiotes* extracts; (F) *m/z* 814.58053 features with highest loading representing the group T6PS.

## 4 DISCUSSION

In this study we opted to test the allelopathic effect of aqueous extracts obtained from the dry biomass of two species of floating macrophytes. We considered that these extracts represent an approximation of a natural condition in a water body, where macrophytes release allelochemicals in the water column, different from that represented by solvent-based extracts such as ethanolic and methanolic ones (X. Wu et al., 2013; Zhou et al., 2014; Lourenção et al., 2021). The aqueous extracts of *P. stratiotes* and *P. crassipes* inhibited the growth of *M. aeruginosa* by different degrees and by different mechanisms.

Using *P. stratiotes* extracts, total inhibition of cyanobacterial growth occurred in 6 days. The phenomenon was previously reported by authors that observed a high inhibitory effect of aqueous extracts of *P. stratiotes* (Lourenção et al., 2021) on *M. aeruginosa* growth. Although in that study, cyanobacterial inhibition occurred with a concentration of 50 mg L^-1^ (here, we used 4.0 g L^-1^), they tested a different methodology to prepare the aqueous extract, which makes comparisons more difficult. Comparing the inhibitory effects of leaf leachates or volatilization, root exudates, and decomposition products of *P. stratiotes* on *M. aeruginosa*, Wu et al. (2015) concluded that root exudation was the main release route for allelochemicals, reinforcing their aqueous soluble nature. In other tests, total inhibition of *M. aeruginosa* growth was achieved using ethanolic extracts or ethyl acetate extracts from *P. stratiotes* (Lourenção et al., 2021; X. Wu et al., 2013).

The *P. crassipes* extract showed a partial (50% - 60%) inhibition of *M. aeruginosa* growth at a concentration of 4.0 g L^-1^, which lasted from day 2^nd^ until day 8^th^. Similar results were reported by Wu et al. (2019), who observed inhibition of *M. aeruginosa* growth using aqueous extract (80 g L^-1^) of the dry biomass of *P. crassipes* roots. De Lima et al. (2023) used exudates from living *P. crassipes* and extracts from decomposing plants (7.5 g L^-1^) to test the inhibition of *Raphidiopsis raciborskii* growth. In this case, extracts from decomposing plants inhibited growth by 60% after 6 days and 100% after 12 days, while exudates from living plants inhibited cyanobacterial growth by a maximum of 25%, indicating that allelochemicals are preserved in the dry biomass and released in aqueous solutions consequently.

In cultures of *M. aeruginosa* exposed to *P. stratiotes*, the increase in growth inhibition over time (from 50% on day 2^nd^ to 90% on day 4^th^ and 100% on day 6^th^) was accompanied by a decrease in Chl-*a* concentrations, almost complete inhibition of the photosynthetic efficiency of PSII from day 4 and lower *psbA* transcript abundance compared to control.

Many allelochemicals from macrophytes, such as phenolic compounds and alkaloids, among others, can inhibit the photosynthetic activity of algae and cyanobacteria (Chen et al., 2021; Hong et al., 2008; Yotsova et al., 2017; Zhao et al., 2020). The underlying mechanism of inhibition was characterized in more detail in studies that tested isolated allelochemicals.

The phenolic molecule N-phenyl-2-naphthylamine (PNA) inhibited *M. aeruginosa* growth after 4 days of exposure (to 0.5 and 1 mg L^-1^), decreased the cellular content of Chl-*a* and other photosynthetic pigments as well as the transcription of photosynthesis-related genes (Qian et al., 2010). In another study (Gao et al., 2017), the repeated exposure of *M. aeruginosa* to low doses of PNA (50 μg L^-1^) resulted in growth inhibition (> 90% in 6 days) and inhibition of PSII. Analyzing the toxicity mechanism of PNA (up to 2 mg L^-1^) in *R. raciborskii* cells, Liu et al. (2015) concluded that the compound decreased electron transport rates through PSII and also observed decreases in cellular Chl-*a* content and growth rate. The investigation of the inhibitory effect of *Pontederia cordata* on *M. aeruginosa* led to the identification of some allelochemicals, including succinic acid (Chen et al., 2021).

Treatment of cyanobacterial cells with succinic acid (60 mg L^-1^) inhibited photosynthesis and, consequently, the growth of *M. aeruginosa*. The mechanism of inhibition of succinic acid involved the damage of the oxygen-evolving complex and reaction center of PSII, blocking photosynthetic electron transport. From this, purified or complex allelochemicals may affect the photosystem structures, simultaneously affecting their functions and potential repair.

Regarding the effect on *psbA* transcripts, the expression of the *psbA* gene increased in the first 12 hours, followed by a downregulation after 24 hours (Chen et al., 2021). The *psbA* gene codes for the D1 protein, a fundamental component of the PSII reaction center involved in the water oxidation reaction complex for subsequent electron transfer (Kojima et al., 2007, p. 200; Li et al., 2016). Once damaged, the D1 protein cannot be repaired, and the cell needs to synthesize new copies (Mulo et al., 2012). Therefore, a corresponding increase in transcripts is expected. In the present study, *psbA* transcript abundance was lower in *M. aeruginosa* cells treated with macrophyte extracts compared to control, possibly the longer period adopted for sampling (from day 2 on) corresponded to the decreased phase in the synthesis of this protein as observed before (Chen et al., 2021).

The generation of reactive oxygen species (ROS) in cyanobacterial cells is an important component of the inhibitory effect of allelochemicals (Zhang et al., 2010; Jing Wang et al., 2011; X. Wu et al., 2015; Ma et al., 2018). Here, ROS concentrations in *M. aeruginosa* cultures were approximately 10 times higher than in the control after 2 days in the presence of macrophyte extracts and increased to 40 times after 4-6 days. It was expected that cyanobacterial cells would increase the expression/activity of antioxidant enzymes in this condition. Accordingly, a higher abundance of *prxA* transcripts was observed 2 days after incubation with the extracts, and this remained until the 4th day for *P. stratiotes* extracts. In contrast, *sod* transcript abundances did not change in the initial time and even decreased later, while SOD activity initially increased in treated cells compared to control.

Similarly, previous studies indicated that *P. stratiotes* extracts cause oxidative stress in target cells. For example, allelochemicals released by *P. stratiotes* roots inhibited the growth of *M. aeruginosa* and increased the cellular content of superoxide anion radicals (X. Wu et al., 2015). Exposure to aqueous and ethanolic extracts of *P. stratiotes* increased the intracellular concentration of hydrogen peroxide in *M. aeruginosa* (Lourenção et al., 2021). Additionally, antioxidant enzyme activities were induced as measured for catalase, peroxidase, and glutathione S-transferase. Conversely, SOD activity declined after exposure to *P. stratiotes* extracts, suggesting that this could be a target for the harmful effect of allelochemicals on cell viability.

Working with isolated allelochemical compounds, Zhang et al. (2010) showed that ρ-coumaric acid and vanillic acid inhibited *M. aeruginosa* growth, and the underlying mechanism was linked to the generation of superoxide anion radicals that caused lipid peroxidation, altering cell membrane permeability and leading to the cyanobacterial cell death. The phenolic compounds catechin and pyrogallic acid, known allelochemicals from submerged macrophytes, induced ROS formation and oxidative stress in *M. aeruginosa* cells (Jing Wang et al., 2011). The allelochemical eugenol causes oxidative stress in *M. aeruginosa* cells mediated by increased intracellular nitric oxide concentration. This was associated with inhibition photosynthetic activity, lipid peroxidation, damage to cell wall and membrane organization, and release of cellular matter (Zhao et al., 2020).

Similarly to our observations, the allelochemical succinic acid induced the expression of the *prxA* gene in *M. aeruginosa* (Chen et al., 2021). This gene encodes the enzyme 2-Cys peroxiredoxin, which reduces peroxide groups using thiol groups as a reducing intermediate, a function related to oxidative stress mediated by both light intensity and chemical imbalance of ROS (Bernroitner et al., 2009). With prolonged exposure, *prxA* gene expression decreased gradually, as also observed for succinic acid (Chen et al., 2021), indicating that oxidative damage exceeded the cellular defense capacity.

Using *P. crassipes* extracts, we observed a different effect on *M aeruginosa* cells. Growth inhibition was limited to a maximum of 60%, Chl-*a* concentrations continued to increase with time. However, less than in the control condition, we did not identify any effect on PSII despite a decrease in *psbA* transcript levels. Exposure to macrophyte extracts led to increase ROS concentrations and a corresponding early but transient increase in *prxA* transcripts. Four days after exposure, cellular growth was constant, and the antioxidant indicators, *prxA* and *sod* transcripts or SOD activity were similar to the control. This indicated that cells could recover from the oxidative condition generated by the extract, probably activating other antioxidant responses that were not assessed in this study. It is also possible that the oxidative damage did not target PSII components, and photosynthesis was not affected, allowing cells to recover growth.

In a co-culture system with *P. crassipes* and *M. aeruginosa,* although the cell biomass of cyanobacteria gradually reduced in 8 days, Chl-*a* content and PSII electron transfer were not affected (Zhou et al., 2014). However, phycocyanin content changed, and pigment ratios were altered, which may have affected the photosynthetic electron transport chain. SOD activity increased transiently and then decreased in comparison to control, while a concomitant increase in malondialdehyde levels indicated lipid peroxidation. Thus, these results support an oxidative condition but limited damage to photosynthetic components of cyanobacterial cells in response to *P. crassipes* allelochemicals. In our study, incubation with the aqueous extract decreased phenolic compounds over time, which may have alleviated the harmful effect on *M. aeruginosa* cells. In contrast, in the co-cultivation system, a continuous release of allelochemical can occur, and the inhibition effect can be more pronounced, as observed by Zhou et al. (2014)

Regarding which chemical compounds would be associated with the effect of macrophyte extracts on cyanobacteria, phenolic compounds can inhibit the growth of cyanobacterial species, and their effects are mediated by oxidative stress (Zhang et al., 2010; Jing Wang et al., 2011; Ni et al., 2013; Ma et al., 2018; He et al., 2023). In the present study, the phenolic concentrations of the aqueous extracts were quantified and, values were higher for *P. stratiotes* than for *P. crassipes*, which coincided with the higher inhibitory effect of the previous macrophyte on *M. aeruginosa* cells, the observed decrease in PSII efficiency and the early induction of *prxA* expression and SOD activity. Upon incubation with *M. aeruginosa* cultures, total phenol concentrations in *P. stratiotes* extracts decreased less than in *P. crassipes* extracts, indicating a prolonged harmful effect on cyanobacterial cells.

For both extracts, total phenol concentrations were reduced over time in the presence of *M. aeruginosa,* indicating the degradation or metabolization of extract compounds due to microbial activity (Neilen et al., 2019). This is supported by the change in the general metabolic profile of the extracts after 6 days, as illustrated by mass spectrometry data. *M. aeruginosa* may have exerted a direct influence on the biotransformation of these aqueous extracts, making them more toxic or, on the contrary, degrading allelochemicals. On the other hand, the composition of extracts incubated for 6 days without cyanobacterial cultures also changed relatively to the initial time. This is due to various factors that can affect these changes, such as temperature, time, pH, chemical composition that might stabilize or destabilize each other, oxygen exposure, and light exposure. For instance, previous studies showed chemical instability of some compounds even at low intensity (30 μmoles.m^-2^ s^-1^) (Jing Wang et al., 2011; Ma et al., 2018; Neilen et al., 2019; Panhota et al., 2023). Bacterial growth in the extracts was tested over these 6 days, and the results were negative (data not shown), ruling out the involvement of bacterial metabolism as the cause of these changes.

It is important to note that *M. aeruginosa* cultures used in this study were non-axenic. Thus, the microbial community associated with the cyanobacterium may have contributed to some results observed as a consequence of incubation with macrophyte extracts. Aqueous extracts would be sources of carbon-based molecules and other compounds that are fundamental to energy acquisition. These potential responses of cyanobacteria-associated microorganisms include ROS generation and possibly SOD activity, which would explain the lack of correspondence between *sod* transcript levels and enzyme activity. Additionally, the microbial community associated with *M. aeruginosa* may have metabolized extract constituents, especially in phenol degradation. The possible participation of this community in allelopathic interactions between macrophytes and cyanobacteria should be further explored.

The practical application of using biomass to inhibit cyanobacterial growth, particularly in this study employing aqueous extracts of macrophytes, presents positive and negative aspects that need to be highlighted. The positive effect is the suppression of cyanobacterial growth, which would make tons of disposable biomass an ecological alternative to water remediation. However, important aspects must be taken into account, such as the release of nutrients from biomass, which could exacerbate the eutrophication in aquatic ecosystems. Additionally, the use of aqueous extract on toxin-producing cyanobacteria blooms should be investigated deeply due to the possibility of cell lysis and, consequently, the release of intracellular compounds. From this, there is a need for monitoring to ensure the overall safety and effectiveness of such interventions, including the potential of cyanobacteria-associated microbiota to degrade these compounds simultaneously. Therefore, although this nature-based approach offers an alternative to mitigating cyanobacterial growth, comprehensive assessments are essential for its implementation in aquatic environments.

## 5 CONCLUSION

Aqueous extracts from dry biomass of *P. stratiotes* and *P. crassipes* can inhibit *M. aeruginosa* growth. Both extracts caused increased ROS production in *M. aeruginosa* cultures, with consequent induction of peroxiredoxin gene expression, but this response did not include a sustained increase in superoxide dismutase activity. The stronger inhibition of the *P. stratiotes* extract on *M. aeruginosa* cells was associated with a decrease in the PSII activity and incapacity of the cells to restore the key protein D1 component, as indicated by a continuous decrease in *psbA* expression. The greater effect of the *P. stratiotes* extract corresponded to a higher concentration of phenolic compounds compared to that of *P. crassipes*. Upon incubation, the macrophyte extracts underwent changes in their chemical composition, including a reduction in the concentration of phenols in the presence of *M. aeruginosa* cultures. The extracts acted differently in reducing *M. aeruginosa* growth. In view of these results, the use of dry macrophyte biomass, which generally accumulates in excess and has no defined destination, can represent an alternative for the sustainable management of freshwater environments impacted by cyanobacterial growth.

## Supporting information

Tabela Suplementar 1

## Acknowledgments

We acknowledge the high-grade PhD fellowship and research grant provided by the Carlos Chagas Filho Foundation for Research Support of the State of Rio de Janeiro (FAPERJ – program POSDOC NOTA 10) for Dr. Allan Santos (E-26/204.609/2021). We also acknowledge students Vitoria Gomes and Julia Marques for their support in the experiments, the Biomolecule Expression, Purification and Analysis Platform (PEPAB-IBCCF-UFRJ), and Dra Carolina Goulart for the use of all equipment necessary to acquire data for this work.

## Supplementary Material

**Table S1.**
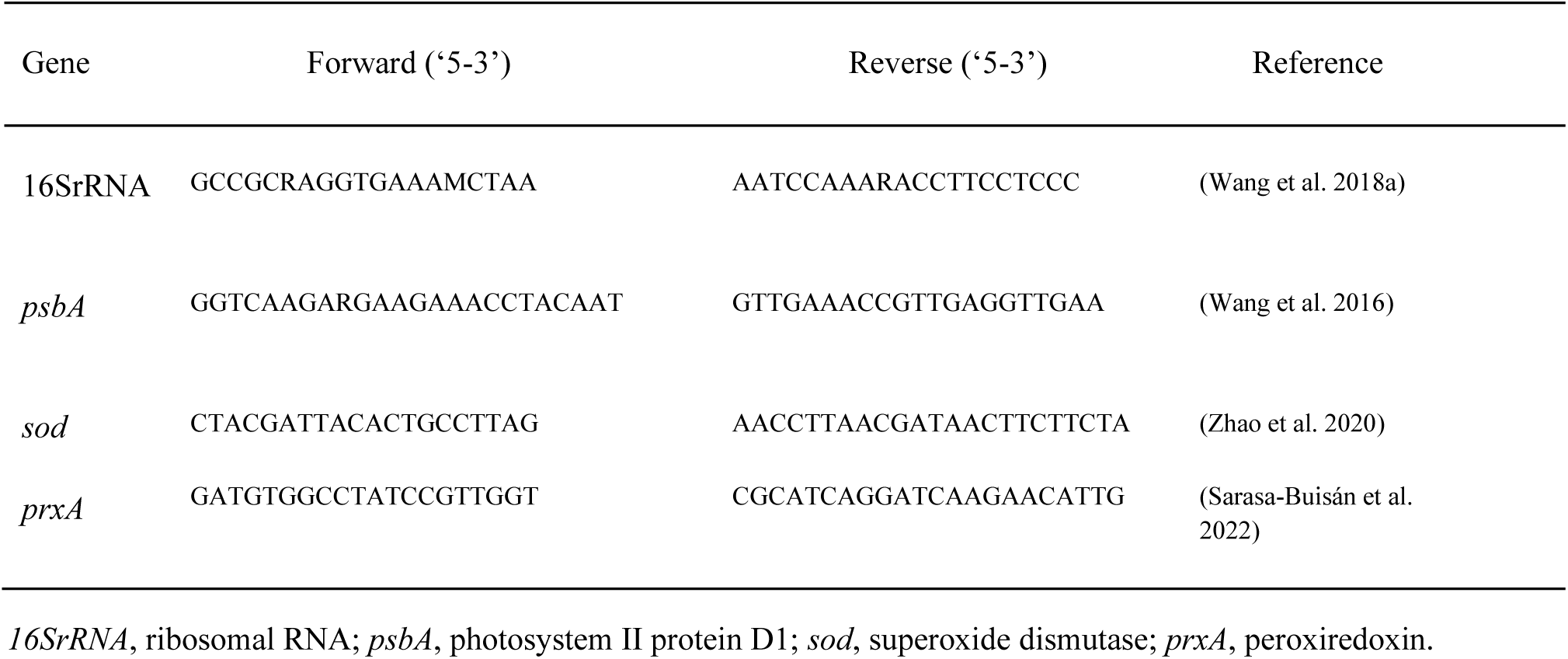
*Primers* used in qPCR.

Negative ionization modes (ESI) data from FT-ICR-MS

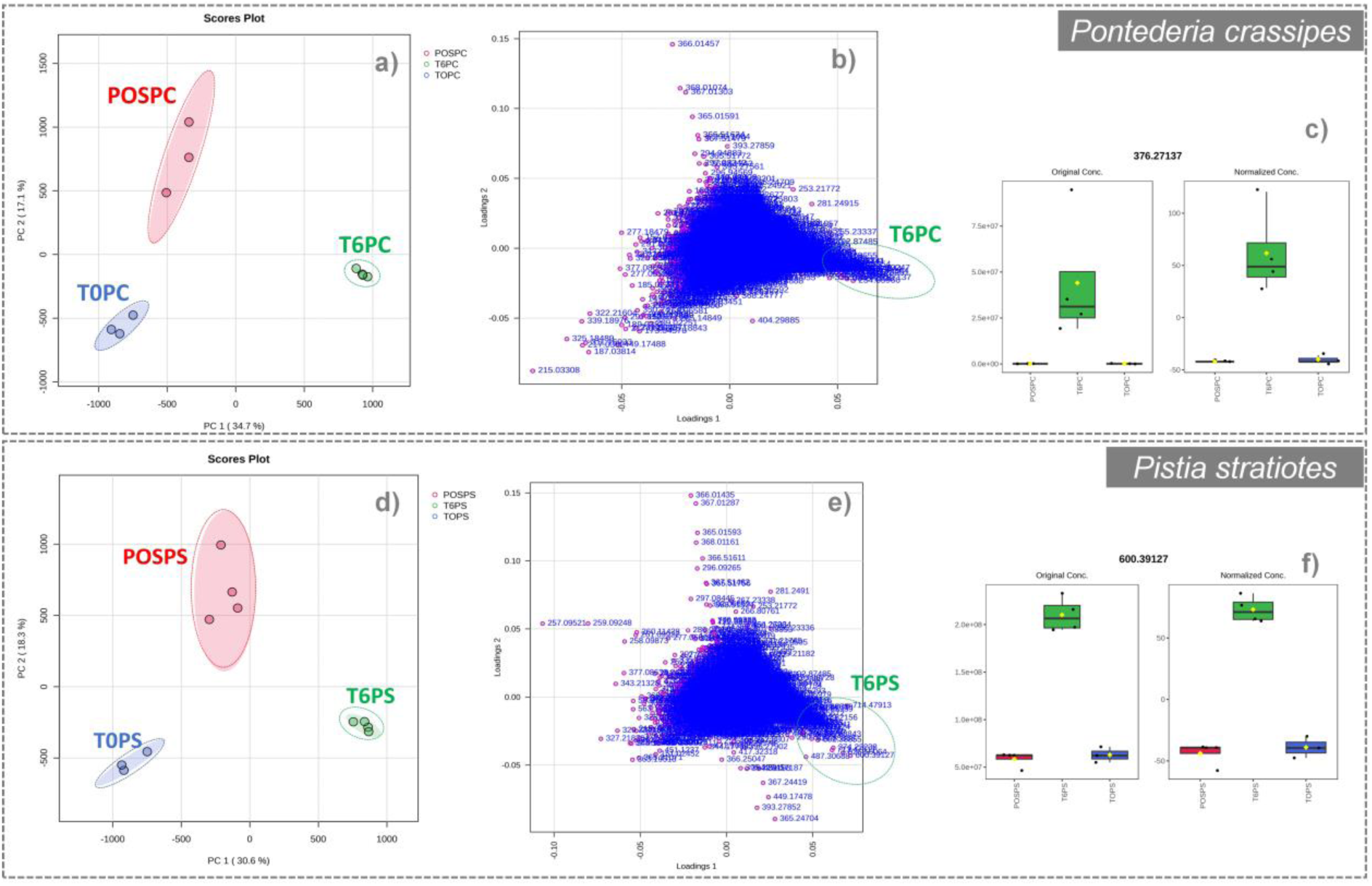

