## Supplementary material for "Physiological response of *Microcystis aeruginosa* exposed to aqueous extracts of *Pistia stratiotes* and *Pontederia crassipes*": Tabela Suplementar 1

**Table S1.** *Primers* used in qPCR.

| Gene | Forward ('5-3') | Reverse ('5-3') | Reference |
|---|---|---|---|
| 16SrRNA | GCCGCRAGGTGAAAMCTAA | AATCCAAARACCTTCCTCCC | (Wang et al. 2018a) |
| *psbA* | GGTCAAGARGAAGAAACCTACAAT | GTTGAAACCGTTGAGGTTGAA | (Wang et al. 2016) |
| *sod* | CTACGATTACACTGCCTTAG | AACCTTAACGATAACTTCTTCTA | (Zhao et al. 2020) |
| *prxA* | GATGTGGCCTATCCGTTGGT | CGCATCAGGATCAAGAACATTG | (Sarasa-Buisán et al. 2022) |

*16SrRNA*, ribosomal RNA; *psbA*, photosystem II protein D1; *sod*, superoxide dismutase; *prxA*, peroxiredoxin.

Negative ionization modes (ESI) data from FT-ICR-MS

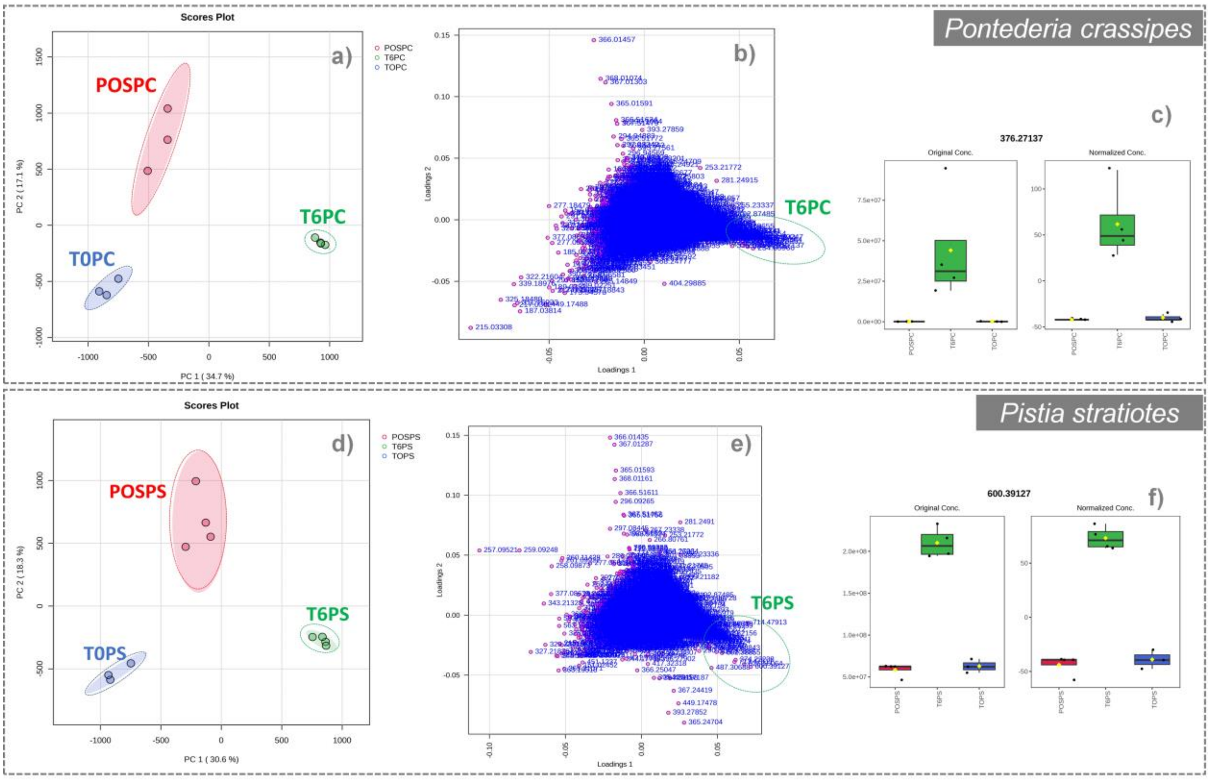
